# Sequence motif dynamics in RNA pools

**DOI:** 10.1101/2024.12.10.627702

**Authors:** Johannes Harth-Kitzerow, Tobias Göppel, Ludwig Burger, Torsten A. Enßlin, Ulrich Gerland

**Affiliations:** Max Planck Institute for Astrophysics Garching, Karl-Schwarzschild-Str. 1, 85741 Garching, Germany; Department of Bioscience, TUM School of Natural Sciences, Technical University Munich, 85748 Garching, Germany; Ludwig-Maximilians-Universität München, Fakultät für Physik, Geschwister-Scholl-Platz 1, 80539 München LMU, Germany; Excellence Cluster Origins, Boltzmannstraße 2, 85748 Garching, Germany

## Abstract

In RNA world scenarios, pools of RNA oligomers form strongly interacting, dynamic systems, which enable molecular evolution. In such pools, RNA oligomers hybridize and dehybridize, ligate and break, ultimately generating longer RNA molecules, which may fold into catalytically active ribozymes. A key process for the elongation of RNA oligomers is templated ligation, which can occur when two RNA strands are adjacently hybridized onto a template strand. Detailed simulations of the dynamics in RNA pools involve a large variety of possible sequences and reactions. Here, we develop a reduced description of these complex dynamics within the space of sequence motifs. We then explore to what extent our reduced description can capture the behavior of detailed simulations that account for the full dynamics in the space of RNA strands. Towards this end, we project the dynamics into a motif space, which accounts only for the abundance of all possible four-nucleotide motifs. A system of ordinary differential equations describes the dynamics of those motifs. Its control parameters are effective rate constants for reactions in motif space, which we obtain from the rate constants for the processes underlying the full dynamics in the space of RNA strands. We find that these reduced motif space dynamics indeed capture important aspects of the informational dynamics of RNA pools in sequence space. This approach could also provide a framework to rationalize and interpret features of the sequence dynamics observed in experimental systems.

## INTRODUCTION

Pools of inter-reacting RNA molecules are a possible starting point for molecular evolution and play a key role in the RNA world hypothesis for the origin of life [4, 11, 20, 21, 24]. The collective behavior of RNA pools, i. e., mixtures of RNA molecules in solution, arises from a complex interplay of intra- and inter-molecular processes. Non-covalent interactions, such as the formation of base-pairs and the stacking of base-pairs into helices, govern the self-folding of RNA strands and the binding between different RNA strands (‘hybridization’ of strands). At the same time, chemical reactions involving covalent bonds govern processes such as template-directed ligation and polymerization, as well as cleavage of RNA strands via hydrolysis. As a consequence, RNA pools are systems with an enormous range of relevant timescales, and a tremendous combinatorial complexity. It is currently neither feasible to track the detailed dynamics of RNA pools experimentally, nor to accurately predict these dynamics computationally. However, the outcome of these dynamics can be assayed experimentally, for instance with mass spectrometry [15] or sequencing approaches [2]. Computational approaches and theoretical analyses can then help to interpret such experiments, and to extrapolate into experimentally inaccessible parameter regimes or to intractable timescales.

In principle, the dynamics of RNA pools could be modeled at various levels of description. However, a very detailed molecular description is computationally intractable when the long-time behavior of a sizable pool is of interest. A hybrid approach like OxRNA [32], which simulates the dynamics of a coarse-grained nucleotide-level RNA model by combining kinetic Monte Carlo and Molecular Dynamics techniques, would be well suited to describe, e. g., the assembly of the RNA complexes required for template-directed ligation. However, this level of description is still too detailed for the slow timescales associated, in particular, with the ligation and cleavage reactions. Prior computational and theoretical work [5, 6, 9, 13, 16, 19, 23, 25–27, 29–31] has therefore used more coarse-grained descriptions that do not explicitly keep track of the polymer conformation of the RNA (or DNA) molecules in the system. Instead, the descriptions for instance keep track of the sequence composition and the strand length distribution of the pool, and estimate the rates for the different kinetic processes included in the model using sequence-dependent base-pairing free energies. For instance, template-directed ligation can occur only when two strands are adjacently hybridized to a template strand. Hence the kinetic rate for this process is proportional to the number of such RNA complexes, which depends on their thermodynamic stability.

While the dynamics of RNA pools is rooted in the polymer physics and chemistry of RNA strands, it can also be regarded from a purely information-centric perspective. For instance, we can consider a string of nucleotides with a specific sequence and count how often it occurs at any position on the RNA strands within a pool. How does the distribution of such sequence motifs evolve in an RNA pool, given a set of intra- and inter-molecular processes? And how do these sequence motif dynamics depend on the rate constants for the different processes? Furthermore, is it possible to describe important features of the sequence motif dynamics in RNA pools with a closed set of dynamical equations, without explicitly describing the underlying physico-chemical processes acting on the RNA strands? These questions motivate the analysis we present here.

As a concrete example and proof of principle, we base our analysis on the ‘RNA reactor’ model and simulation framework [9, 13, 23], which models the interplay of four processes: RNA hybridization, de-hybridization, cleavage, and template-directed ligation. Focusing on the case of 4-nucleotide motifs, we devise motif rate equations that describe the time evolution of motif concentrations under the simultaneous action of the four physico-chemical processes of the RNA reactor. Our approach can be regarded as a generalization of the 2-nucleotide motif dynamics of Ref. [26]. Importantly, for our 4-nucleotide motif dynamics, we can establish a direct relation between its kinetic parameters and those of the underlying physico-chemical processes acting on the RNA strands. Four is the minimal motif length, for which this is possible, since the RNA reactor model includes a kinetic effect in ligation that depends not only on the identity of the nucleotides at the ligation site, but also on the neighboring nucleotides to both sides. This kinetic stalling of template-directed ligation after mismatches has been established experimentally [14, 22] and can have a significant impact even on the qualitative behavior of RNA pools [9].

The mapping between the kinetic parameters of the full RNA reactor model and those of the motif rate equations permits us to test which aspects of the dynamics are retained by the reduced motif description, and which are lost. Towards this end, we consider each of the five different scenarios of Ref. [9] for the RNA reactor also within the motif rate equations. In the following Methods section, we therefore first briefly summarize the RNA reactor model before developing the reduced description in motif space. The comparison between the two levels of description then follows in the Results section.

## METHODS

In this section, we introduce motif rate equations as chemical rate equations for motif concentrations, and determine their parameters from those of the strand-based ‘RNA reactor’ model [9, 13, 23], see Fig. 1 for illustration. The ‘RNA reactor’ model describes the kinetics of oligonucleotide strands that can take on different configurations in linear (non-branched) complexes, but do not fold onto themselves. These kinetics are simulated using the Gillespie algorithm [7, 8], with kinetic rate constants that are chosen consistent with the thermodynamics of hybridization between the oligonucleotide strands (in the following, Δ𝒢_tot_ denotes the total free energy difference between a configuration where a single strand is bound in a complex and the configuration where this strand is unbound). Here, we specifically use the energy model of Ref. [9] for binary nucleotide alphabets, which allows for an energy bias for alternating sequences, such that fully hybridized blocks of two by two nucleotides have a hybridization energy difference Δ*γ* if they consist of perfectly matching alternating 2-nucleotide motifs compared to perfectly matching homogeneous 2-nucleotide motifs. (If the configuration involves a dangling end, i. e., a 2-nucleotide motif is hybridized to the end of a strand such that one nucleotide has no binding partner, the corresponding energy bias for alternating sequences is denoted by *δ*_*ϵ*_.) Furthermore, single strands can be cleaved at any bond with a rate constant *k*_cut_, independent of the nucleotide sequence context, and templated ligation is characterized by a ligation rate constant *k*_lig_. In the presence of mismatches (non-complementary hybridization of nucleotides) directly at the ligation site, the ligation rate constant is reduced by a stalling factor *σ*_1_, while next-nearest neighbor mismatches stall ligation with the factor *σ*_2_. We consider a closed system without influx and outflux of strands or nucleotides.

**FIG. 1.**
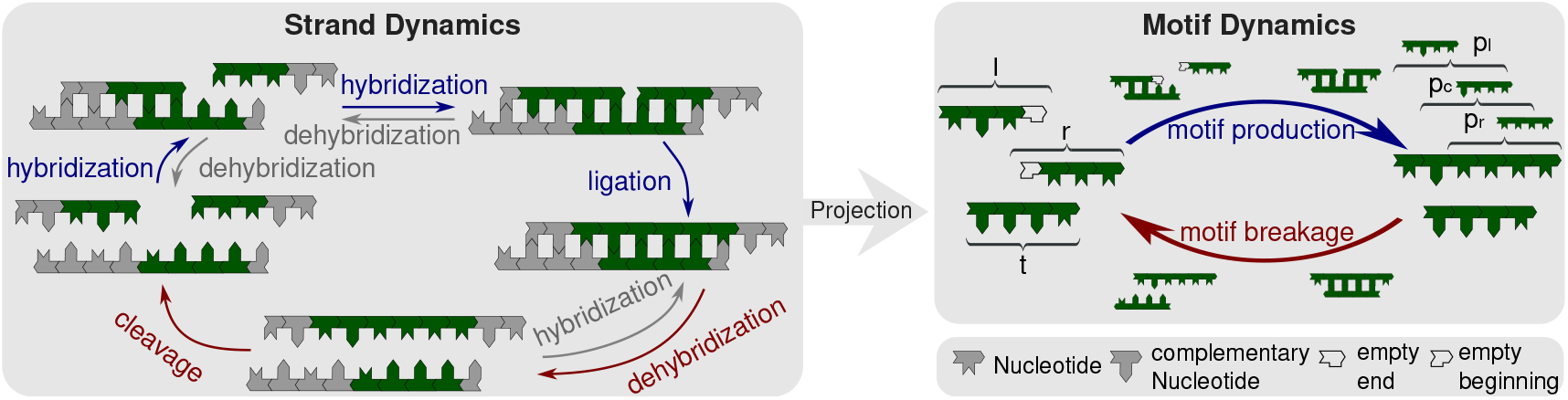
Kinetic processes of the strand-based ‘RNA reactor’ model [9, 13, 23] and their projection onto motif kinetics. We consider heteropolymer strands consisting of the two complementary nucleotides X and Y, which could represent RNA or DNA nucleotides. (a) Schematic of the reactions within the strand reactor, including hybridization, dehybridization, templated ligation, and cleavage of oligonucleotide strands. Strands that are not covalently linked are separated by small gaps, and four-nucleotide motifs, including 3^*′*^-or 5^*′*^-ends, are highlighted in green. In templated ligation, a 3^*′*^- and a 5^*′*^-end motif build a 6-nucleotide motif that contains three overlapping 4-nucleotide motifs. (b) At the level of sequence motifs, the kinetics are based on motif production and motif breakage reactions. In four-nucleotide motifs, strand ends are marked with 3^*′*^ or 5^*′*^ respectively. Templated ligation generates longer strands containing additional motifs, thereby contributing to motif production, while strand cleavage leads to to motif breakage. Hybridization is not explicitly represented in the motif-level description; instead, it is implicitly incorporated into the rate constants for motif production and breakage by assuming a hybridization-dehybridization equilibrium and deducing the corresponding dissociation constants.

### Motif concentrations

Before we introduce the motif rate equations, we specify how we track the motif concentrations, which we group into a motif concentration vector 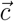. Towards that end, we define the motifs we track, introducing conventions to prevent double counting of motifs, and present how we convert strand numbers into motif concentrations.

To establish a proof of principle for our approach, we use a minimal binary nucleotide alphabet 𝒜 = *{*X, Y*}*, consisting of the two generic complementary nucleotides X and Y. These could represent, e.g., the RNA nucleotides A and U, or the DNA nucleotides A and T. Based on this alphabet, we consider sequence motifs that are directed (chemically from the 5’ to the 3’ end of a strand). To track beginnings and ends of strands, we introduce the additional symbol 0, and denote with 𝒜_0_ = 𝒜 ∪ *{*0*}* the alphabet including 0. Since the ligation rate constant in the strand reactor simulation depends on the two nucleotides to the left and the two nucleotides to the right of the ligation site, the minimal motif length required to project the strand kinetics on the motif space is four. We thus choose to focus on the 4-nucleotide motif dynamics. We track monomers and dimers explicitly, while all longer strands are tracked by their containing 4-nucleotide motifs, including their beginning and end motifs, i. e. motifs that start, or respectively end, with an empty spot symbol 0, but contain only letters elsewhere (Fig. 1b). We refer to beginning motifs simply as beginnings, end motifs as ends, and call 4-nucleotide motifs that do not contain a 0 but only letters continuations.

We avoid double counting of motifs by introducing the convention that the second spot of a 4-nucleotide motif is always a letter. The first and the last spot can be a letter or empty. This allows for ends, beginnings, and continuations, as well as for monomers and dimers. The third spot is only empty for monomers such that empty spots do not occur in between letters. To determine motif concentrations from a given state of the strand reactor, we define the concentration of a single particle, 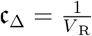, i. e., the concentration of one complex in the strand reactor with a volume *V* _R_. The motif concentrations are then obtained by multiplying c_Δ_ with the counts of all 4-nucleotide motifs in the strands of the reactor (following the above conventions).

Note that our description of the motif dynamics in terms of motif rate equations (to be introduced below) corresponds to a mean field approach, which implicitly assumes the thermodynamic limit of the reactor volume *V* _R_ going to infinity at constant concentrations, such that the dynamics can be described by chemical rate equations instead of a chemical master equation. To nevertheless take finite size effects into account, we introduce a lower threshold for reaction rates, such that reations cannot take place if any of its substrates have a concentration below a certain threshold (see Appendix A 4 for more details and Refs. [1, 3, 12, 28] for similar approaches).

### Motif rate equations

We now turn to the construction of the motif rate equations for a closed reactor, without influx or outflux. We consider the limit, in which ligation is much slower than hybridization and dehybridization, such that a hybridization-dehybridization equilibrium is reached, see Table I for the involved parameters. The fraction of single-stranded versus hybridized strands is then determined by dissociation constants,

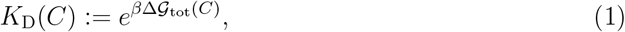

for each complex configuration *C*. In this limit, we incorporate templated ligation together with the dissociation constants in the motif production terms, 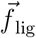. Analogously, we derive motif breakage terms, 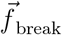, from the dissociation constants and strand cleavage. Thus, the total motif rate equation contains these two classes of reaction terms:

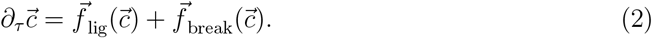

**TABLE I.** Hybridization parameters of the motif rate equations and the strand reactor simulation.

| Symbol | Parameter |
| --- | --- |
| $\bar{\gamma}_{\text{com}}$ | average energy value of complementary (nearest neighbor) blocks |
| $\gamma_{1\text{nc}}$ | stacking energy for blocks with one non-complementary nucleotide pair |
| $\gamma_{2\text{nc}}$ | stacking energy for blocks with two non-complementary nucleotide pairs |
| $\Delta\gamma$ | stacking energy difference (hybridization bias) of alternating and homogeneous blocks |
| $l_{\text{low}}$ | lower bound dehybridization length |
| $\epsilon_{\text{com}}$ | average energy value of complementary blocks with dangling end |
| $\epsilon_{1\text{nc}}$ | stacking energy for blocks with dangling end and one non-complementary nucleotide pair |
| $\delta_{\epsilon}$ | stacking energy difference of alternating and homogeneous blocks with dangling end |
| $\Delta\mathcal{G}_{\text{hyb}}$ | hybridization Gibbs free energy |
| $k_{\text{on}}$ | hybridization rate constant |
| $k_{\text{off}}$ | dehybridization rate constant |
| $K_{\text{D}}$ | dissociation constant |
| $\tilde{K}_{\text{D}}$ | effective dissociation constant |

Motif production, the first set of terms in the motif rate equation captures hybridization and dehybridization implicitly and ligation explicitly, see Table II for the involved parameters. The corresponding motif production rate constants are denoted 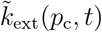, with produced motif *p*_c_ onto the templating motif *t*.

**TABLE II.** Ligation parameters of the motif rate equations and the strand reactor simulation.

| Symbol | Parameter |
| --- | --- |
| $k_{\text{lig}0}$ | default ligation rate constant |
| $\sigma_i$ | stalling parameter, $i \in \{1, 2\}$ |
| $\Phi_{\pm}$ | stalling factors |
| $k_{\text{lig}}$ | ligation rate constant |
| $k_{\text{ext}}$ | extension rate constant |
| $\tilde{k}_{\text{ext}}$ | motif production rate constants |

Motif production describes the process in which two terminal strands hybridize adjacent to one another on a template strand, see Figure 1. The left strand involved in the ligation carries the terminal reactant motif *l* with an open 3’ end, *l* = (*l*_1_, *l*_2_, *l*_3_, 0) (*l*_1_, *l*_2_, *l*_3_ ∈ 𝒜). For a monomeric left strand, *l* = (0, *l*_2_, 0, 0) (*l*_1_, *l*_3_ = 0); for a dimer, *l* = (0, *l*_2_, *l*_3_, 0). The right strand contains the beginning reactant motif *r* with an open 5’ end, *r* = (0, *r*_2_, *r*_3_, *r*_4_) (*r*_2_ ∈ 𝒜, *r*_3_, *r*_4_ ∈ 𝒜_0_), where *r*_4_ = 0 and *r*_3_ ∈ 𝒜 for dimers, and *r*_3_ = 0 = *r*_4_ for monomers.

The template contains the motif *t* = (*t*_1_, *t*_2_, *t*_3_, *t*_4_) directly at the ligation site. Note that in our model the template sequence does not need to be fully complementary to the product motif. However, depending on the energy parameters, hybridization and templated ligation may stall in the presence of mismatches.

Then the two strands with motifs *l* and *rrm* ligate to the product motifs

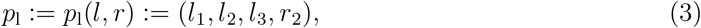

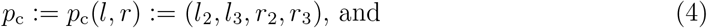

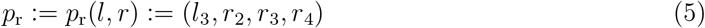

and eventually dehybridize again. In case the left (right) reactant is a monomer, there is no first produced motif *p*_l_ (third produced motif *p*_r_) and the central and right (left) produced motif become *p*_c_ = (0, *l*_2_, *r*_2_, *r*_3_) and *p*_r_ = (*l*_2_, *r*_2_, *r*_3_, *r*_4_) (*p*_l_ = (*l*_1_, *l*_2_, *l*_3_, *r*_2_) and *p*_c_ = (*l*_2_, *l*_3_, *r*_2_, 0)). In case both reactants are monomers, there is only one produced motif *p*_c_ = (0, *l*_2_, *r*_2_, 0).

Since we only consider nearest neighbour contributions to the effective ligation rate, 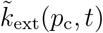 does neither depend on the first nucleotide of the ending strand (*l*_1_) nor the last nucleotide of the leaving strand (*r*_4_). The reaction takes place with the total motif production rate Λ_*l,r,t*_,

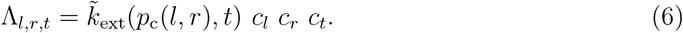

Details of its calculation are in Appendix A 1. With this, the motif production terms split up into five sets of terms. Two for the ending and beginning reactant, 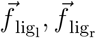 and three terms for the three produced motifs, collected in 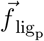.

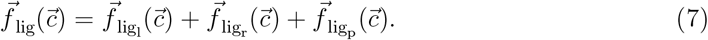

Both, the ending as well as the leaving reactant, are consumed in this reaction. This leads to a negative contribution in the motif rate equation,

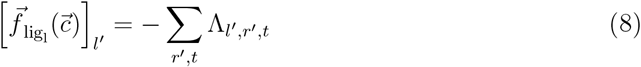

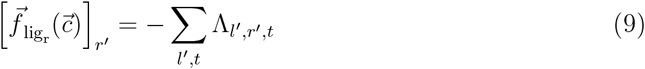

for ends, i. e., possible ending reactants including monomers, and beginnings, i. e., possible beginning reactants. For all other motifs, 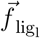 and 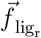 become zero.

For the three produced motifs (*p*_l_, *p*_c_, *p*_r_), we get positive contributions in the motif rate equation,

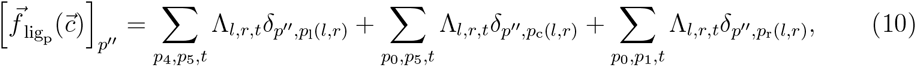

without a contribution of any other motif. For convenience, we chose a compact notation without stating the summation ranges explicitly. The ranges following from our convention are discussed in detail in Appendix A 1.

By splitting the motif production terms into five subterms for the two reactants and three motifs, we have set up the motif production terms of the motif rate equation. For the motif production rate constants, we use the extension rate constants derived in [9] and discussed in Appendix A 2 for complexes up to length four. Breakage leads to two terms in the motif rate equation. First, a negative term 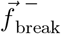 for the motif that breaks and, second, a positive term 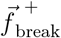 for motifs that result from breakage. Before we discuss the terms for the motif rate equation, we draw the connection to the strand reactor simulation to derive the breakage rates from cleavage rate constant and dissociation constant, see Table III. Breakage of motifs is the reaction that models the effect of cleavage in the strand reactor simulation. For the motif rate equations, we have to adjust the rate constant by taking the possible protection due to hybridization into account. In the strand reactor simulation, cleavage is modeled by a sequence independent cleavage rate constant *k*_cut_ for any single stranded segment. Since we do not store hybridizations in the motif dynamics, we need to reconstruct their occupation using the dissociation constant which gives the fraction of free versus occupied strands as described above.

**TABLE III.** Cleavage parameters of the strand reactor simulation and breakage parameters of the motif rate equations.

| Symbol | Parameter |
| --- | --- |
| $k_{\text{cut}}$ | cleavage rate constant |
| $\tilde{k}_{\text{cut}}$ | effective cleavage rate constant |
| $\phi_{-}$ | left continuation frequency |
| $\phi_{+}$ | right continuation frequency |
| $\tilde{k}_{\text{cut}}$ | breakage rate constant |

In principle, there are three breakage spots in every 4-nucleotide motif. Since we lose correlations longer than the motif length, we approximate the stability of a hybridization site by the dissociation constant for a single 4-nucleotide motif interpreted as a tetramer, i. e. assuming it to be a strand of four nucleotides only, as we did for the motif production rate constants. In fact, hybridization can be stronger, if the overlap is actually longer, which consequently results in an underestimation of the number of double stranded segments in the motif rate equations. We obtain the breakage rate by multiplying the breakage rate constant with the dissociation constant. Since the fraction of double stranded segments is underestimated, the breakage rate constant is thus overestimated in the motif rate equatitons. We leave the introduction of a correction term for this for future work.

Now that we have clarified the connection to cleavage, we can calculate the breakage rate constants. In a 4-nucleotide motif there are three breakage spots that we denote by the index *i* ∈ *{*1, 2, 3*}*. For the central breakage spot (*i* = 2), we use the effective dissociation constant 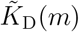, i. e. the dissociation constant of the strand *m* = (*m*_1_, *m*_2_, *m*_3_, *m*_4_) for all configurations in which the central bond (*m*_2_, *m*_3_) is hybridized to any other motif. In the case of similar concentrations of all hybridization partners (templates), it is calculated analogously to the sequence-averaged dissociation constant in [9]: the inverse average of inverse dissociation constants of the single configurations *C* = [*m, m*^*′*^], but by summing over the template for a given sequence. In case of the effective dissociation constants those are the hybridizations onto different templates *m*^*′*^.

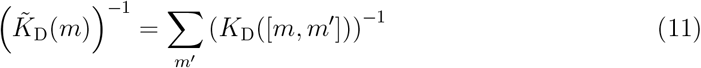

Note that the assumption of similar concentrations for all hybridization partners leads to an overestimation of the hybridization rate for motifs that are actually less abundant, and to an underestimation for those present at higher concentrations. The effective dissociation constant is therefore an approximation that underestimates the dissociation for low-concentration motifs and, overestimates it for highly concentrated motifs. Moreover, longer hybridization sites are reduced to motif length, which causes the hybridization rates of longer strands to be underestimated and their effective dissociation constant to be overestimated.

Then the effective cleavage rate constant 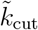 is given by the cleavage rate constant *k*_cut_ multiplied by the effective dissociation constant.

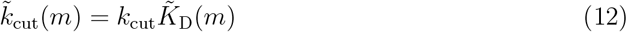

We approximate the (central) motif breakage rate constant 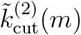 by the effective cleavage rate constant 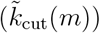. A central breakage also implies breakage of the right bond of the preceeding motif and the left bond of the subsequent motif. The corresponding left- and right-bond breakage rate constants must therefore be determined consistently from the central breakage. To this end, we extend the left (right) breaking motif to the left (right) according to the left (right) continuation frequency *ϕ*_−_ (*ϕ*_+_), defined as the concentration of the specific continuation divided by the sum of the concentrations of all possible continuations, including a beginning (end),

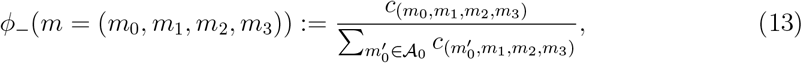

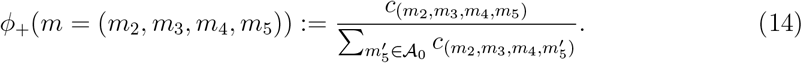

Then we marginalize over the continuations to the left (right) and corresponding breakage rate constants to get effective breakage rate constants 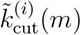 for the left (*i* = 1), central (*i* = 2) and right (*i* = 3) bond [33],

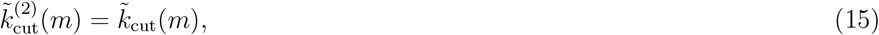

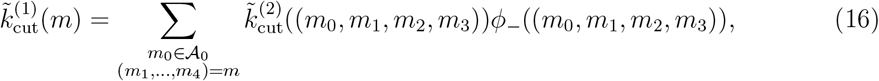

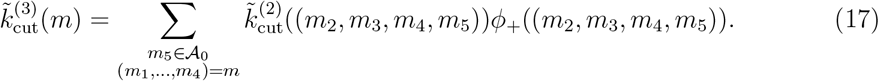

Given the effective breakage rate constants, we can now set up the rates for the breaking motifs 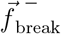 and the resulting ones 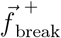

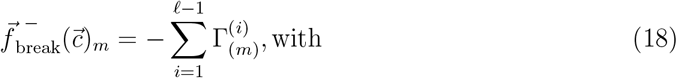

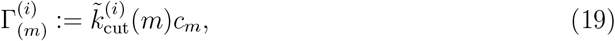

and

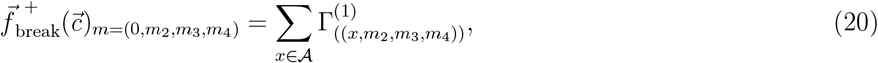

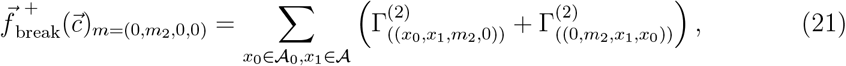

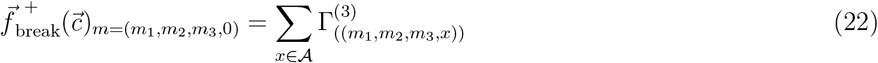

with *m*_2_, *m*_3_ ∈ 𝒜 and *m*_1_, *m*_4_ ∈ 𝒜_0_. The central breakage term (21) only produces motifs in case they are monomers, as all other broken motifs are already considered in the left and right breakage terms.

In summary, the total motif rate equation is given by the sum of motif production terms and breakage terms. Hybridization and dehybridization are captured in all rates implicitly by approximating hybridization-dehybridization equilibrium via dissociation constants. Approximating motifs by strands of the same length as the motifs, we calculated corresponding motif production rate constants and breakage rate constants from the parameters of the strand simulation. For details on its implementation, see Appendix A.

## RESULTS AND DISCUSSION

To test and validate our projection of the strand dynamics onto motif rate equations, we compare the respective dynamics for five different parameter sets. We directly compare the motif concentration trajectories, but also use observables such as motif entropy and mean strand length, where the latter can be reconstructed from the motif concentrations as we will show. Before discussing the results, we describe the five different parameter sets and our choice of initial conditions.

### Parameter Sets and Initial Conditions

In order to compare the sequence motif dynamics in the strand reactor simulation to the solutions of the motif rate equations, we use five parameter sets labeled 0 to 4, which focus on different combinations of three features described above [9]: ligation stalling (s), cleavage (c), and hybridization bias (h). For each feature, we use two parameter values, which either essentially deactivate or activate the corresponding feature, e.g., (*¬*s, s) for ligation stalling, see Table IV. Parameter set 0 deactivates all three features (*¬*s, *¬*c, *¬*h). Parameter set 1 has ligation stalling activated (s, *¬*c, *¬*h), while parameter set 2 additionally activates cleavage (s, c, *¬*h). In parameter set 3, cleavage is deactivated again, but hybridization bias is activated (s, *¬*c, h), and finally in parameter set 4, all three features are activated (s, c, h). The remaining parameters that are kept the same in all five scenarios are discussed in Appendix A 2 and stated in Table V.

**TABLE IV.** Parameter sets used for the strand reactor simulations. The top table shows, for each varied parameter, the two values referred to as ‘0’ and ‘1’ below. The bottom table lists the chosen value combinations for the five parameter sets.

| Parameter |  | Value 0 | Value 1 |
| --- | --- | --- | --- |
| Ligation Stalling | $\sigma_1$ | 1.0 | 0.05 |
| | $\sigma_2$ | 1.0 | 1.0 |
| Cleavage | $k_{\text{cut}}$ | $4.24 \cdot 10^{-18}$ | $9.056 \cdot 10^{-11}$ |
| Hybridization Bias | $\Delta\gamma$ | 0 | -0.3 |
| | $\delta_\epsilon$ | 0 | -0.15 |
| Parameter Set | Ligation Stalling | Cleavage | Hybridization Bias |
| 0 | 0 | 0 | 0 |
| 1 | 1 | 0 | 0 |
| 2 | 1 | 1 | 0 |
| 3 | 1 | 0 | 1 |
| 4 | 1 | 1 | 1 |

**TABLE V.** Reaction parameters of the strand reactor simulation.

| Parameter | $k_{\text{lig0}}$ | $\epsilon_{\text{com}}$ | $\epsilon_{1\text{nc}}$ | $\bar{\gamma}_{\text{com}}$ | $\gamma_{1\text{nc}}$ | $\gamma_{2\text{nc}}$ |
| --- | --- | --- | --- | --- | --- | --- |
| Value | $3.73 \cdot 10^{-6}$ | -0.625 | 0.375 | -1.25 | 0.375 | 0.75 |

In most cases, we use the same initial conditions as in Ref. [9]: The closed RNA reactor contains a total mass of 5000 nucleotides, initially predominantly as monomers (2460 of each nucleotide, X and Y) in an unbiased mix with a few dimers (10 dimers of each kind, XX, XY, YX, YY). The reactor volume is such that the concentration of a single molecule is c_Δ_ = 2 *·* 10^−6^c_0_, where 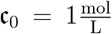 denotes the reference concentration. In the case of parameter set 2, we also probe the dependence of the dynamics on the initial conditions, and thus vary the initial conditions away from these default values. For the motif rate equations, we convert the initial conditions into the corresponding motif concentrations, as described above, i.e., we scan each strand for all its containing 4-nucleotide motifs and multiply the resulting total motif counts with c_Δ_.

### Templated Ligation without Cleavage, Ligation Stalling and Hybridization Bias

We first consider the case of negligible cleavage, without ligation stalling nor hybridization bias (parameter set 0). In this scenario, templated ligation only favors the extension of complementary strands because of hybridization energy differences, but not stalling in the ligation itself. Fig. 2 illustrates the resulting motif dynamics by showing motif concentration trajectories for X monomers, YY dimers, XYX beginnings, and XXXX motifs (to capture the behavior on many scales, both the concentration and the time axis have a logarithmic scale). Here, the solution of the motif rate equations (green dashed lines) is superimposed on an ensemble of stochastic trajectories from the strand reactor simulation.

**FIG. 2.**
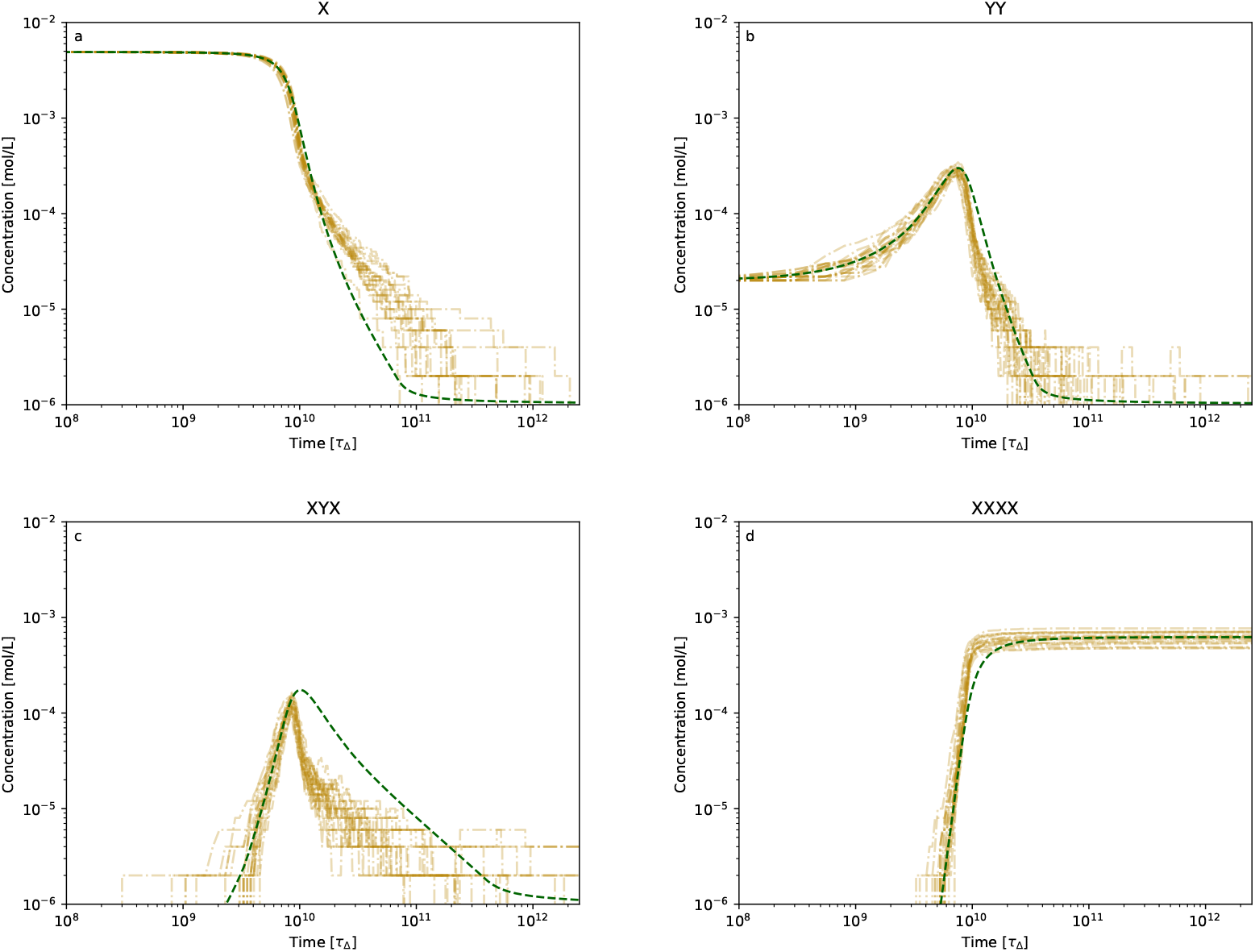
Motif concentrations in the strand reactor simulation (gold) and motif rate equations (green) of the motifs as stated for parameter set 0, i. e.without hybridization bias, without ligation stalling and negligible cleavage.

The trajectories in Fig. 2 reflect the elongation dynamics of strands via templated ligation. Monomers, which are initially dominant in the pool are depleted, generating dimers and longer strands. The dimer concentrations first rise from their small initial values, then reach a peak, before dimers are depleted together with the monomers. The 3-nucleotide beginnings (trajectory for XYX) display a concentration peak shortly after the dimers, with a concurrent rise of the 4-nucleotide motifs, which then reach a stationary value. The elongation dynamics is directly seen in Fig. 3, which shows the mean strand length *L* as a function of time. Here, the mean-field approximation to the mean strand length (green dashed line) is computed from the motif rate equations as described in Appendix C 1.

**FIG. 3.**
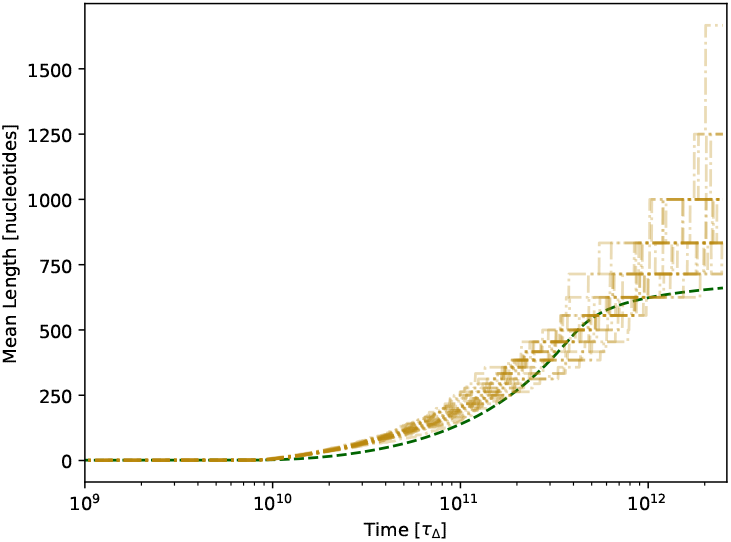
Mean length of the strand reactor simulation (dash dotted gold, mean solid gold) and the motif rate equations (dashed green) for parameter set 0 without hybridization bias, without ligation stalling and negligible cleavage.

As shown in Figs. 2 and 3, the motif rate equations capture many key aspects of the stochastic strand dynamics in the RNA reactor for parameter set 0. During the initial phase, when the mean strand length is shorter than the motif length, the agreement is quantitative. At later times, however, elongation in the motif rate equations lags behind the strand reactor simulation. This lag arises because the motif-level description cannot account for the slow dehybridizatoin of strands longer than the four-nucleotide motif length. For strands with more than four matching hybridization bonds, the hybridization energy is underestimated, causing the motif rate equations to overestimate the dissociation constants. As a consequence, the extension rate constants for longer strands are underestimated. Conversely, when long strands in the strand reactor become trapped in unproductive double-stranded configurations, they can no longer serve as templates. In the motif rate equations such blocked states either do not occur or, if they do, the strands dehybridize more rapicly due to the imposed maximum hybridization energy of four perfect matches. The interplay of these effects has been examined in detail in Ref. [23]. Notably, the elongation behavior in the motif rate equations eventually catches up with the strand reactor simulation at long times. This is partly because the motif rate equations describe the thermodynamic limit of an infinite number of particles, allowing strands to grow without bound, whereas in the strand reactor simulationgrowth is constrained by the fixed total mass of 5000 nucleotides in the finite RNA reactor (see Fig. 3). In principle, the finite size of the RNA pool could influence the dynamics in the strand reactor already at the onset of growth. However, in our case the pool is sufficiently large to avoid such an effect, as estimated in Appendix C 2.

Besides the elongation dynamics, the dynamics of the RNA reactor in sequence space is of key interest to characterize its behavior. Towards this end, Ref. [9] introduced the “zebraness” *Z* as a system-level observable measuring an asymmetry that can spontaneously arise in an RNA reactor. The system-level zebraness is defined as the fraction of alternating 2-mers, ‘XY’ and ‘YX’ within the sequence pool. Hence *Z* takes values between 0 and 1.

If the pool contains only homogeneous motifs (‘XX’, ‘XX’), the zebraness is 0, while it is 1, if the pool contains only zebra-like motifs (‘XY’, ‘YX’). If the pool is unbiased and all motifs occur with the same frequency, the zebraness is 0.5. In terms of 4-nucleotide motifs, the system-level zebraness is calculated via

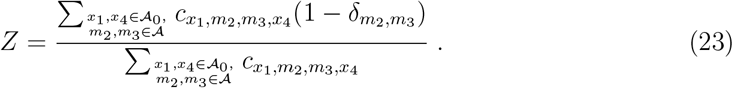

Another relevant observable for the distribution in sequence space is the motif entropy.

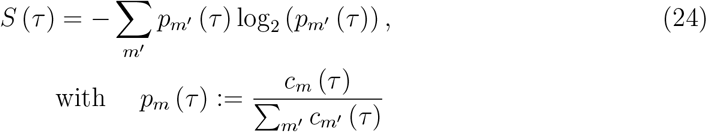

for all motifs *m* (and *m*^*′*^), including monomers, dimers, beginnings, continuations and ends, which are considered in the sums. The dynamics of both observables, the zebraness *Z* and the motif entropy, are shown in Fig. 4 for parameter set 0. Since in this case there is no hybridization energy bias for alternating motifs, the system-level zebraness *Z* stays at its initial level of 0.5 for the motif rate equations (Fig. 4a, dashed green line). This is also the ensemble average of the strand reactor simulation trajectories, which experience some stochastic fluctuations at the onset of growth, at which ligations of longer strands become more frequent than ligations of two monomers.

**FIG. 4.**
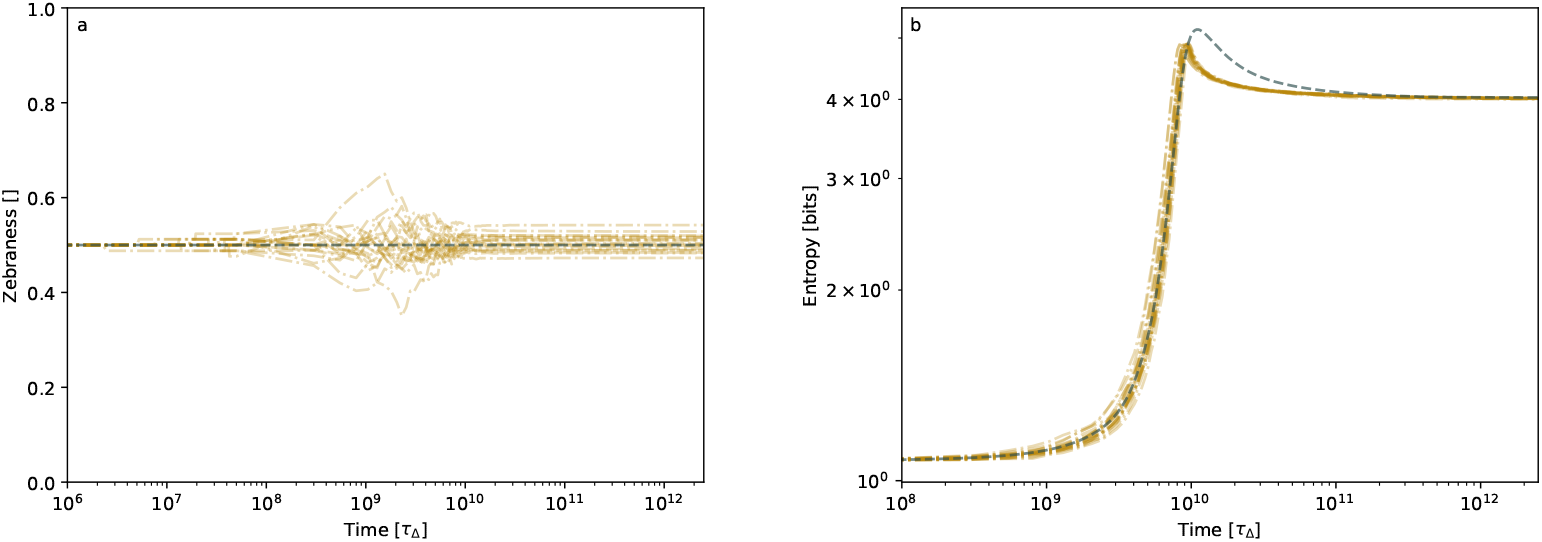
System-level zebraness (a) and motif entropy (b) of the strand reactor simulation (dash dotted gold, mean solid gold) and the motif rate equations (dashed green) for parameter set 0 qualitatively agree.

In contrast, the motif entropy shows a distinctive feature in its dynamics (Fig. 4b), recapitulating the behavior already observed above for the individual motifs (Fig. 2). After a growth phase, the entropy displays a transient maximum, and then relaxes to a stationary value. The motif rate equations quantitatively describe the growth phase, lag behind the strand reactor simulation during the relaxation phase, but ultimately reach the correct stationary value of slightly more than 4 bits. This is the entropy of evenly distributed continuations only, with two beginning and two ending motifs, after all other beginnings and ends have been extended via templated ligation.

Taken together, most aspects of the strand dynamics are captured well by the motif rate equations for parameter set 0. Expected deviations due to stronger hybridization are present, but do not affect the qualitative behavior of the dynamics and are also quantitatively small. Obviously, solving the motif rate equations amounts to a much lower computational effort than the strand reactor simulation (by about four orders of magnitude for the present case). We next explore whether these conclusions remain true also in the presence of cleavage, ligation stalling, and hybridization bias. Since we found that for the sake of comparing the motif rate equations to the strand reactor simulation, the motif entropy recapitulates the behavior of the individual motif concentration trajectories, we will focus on the motif entropy as an aggregate observable for the remaining parameter sets.

### Ligation Stalling, Cleavage and Hybridization Bias

We first consider the effect of ligation stalling (parameter set 1), which slows down templated ligation in the presence of mismatches close to the ligation site. Accordingly, the time evolution of the mean strand length, shown in Fig. 5a, is slightly slower compared to the case without ligation stalling (Fig. 3), but otherwise not significantly altered. The same holds true for the time evolution of the motif entropy (Fig. 6b) compared to the case without ligation stalling (Fig. 4b). For both observables, the motif rate equations capture the behavior of the strand reactor simulation to the same degree as for parameter set 0.

**FIG. 5.**
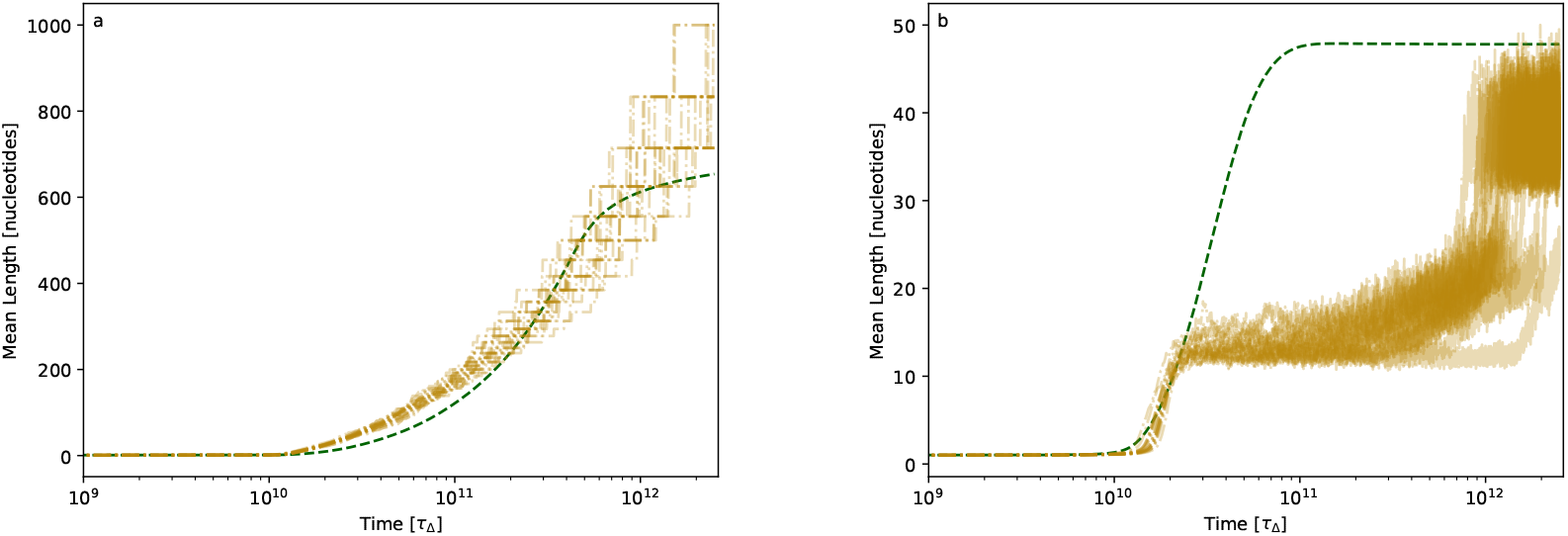
Time evolution of the mean strand length, as in Fig. 3, but for (a) parameter set 1 and (b) parameter set 2, where, in the latter, ligation stalling and cleavage are activated.

**FIG. 6.**
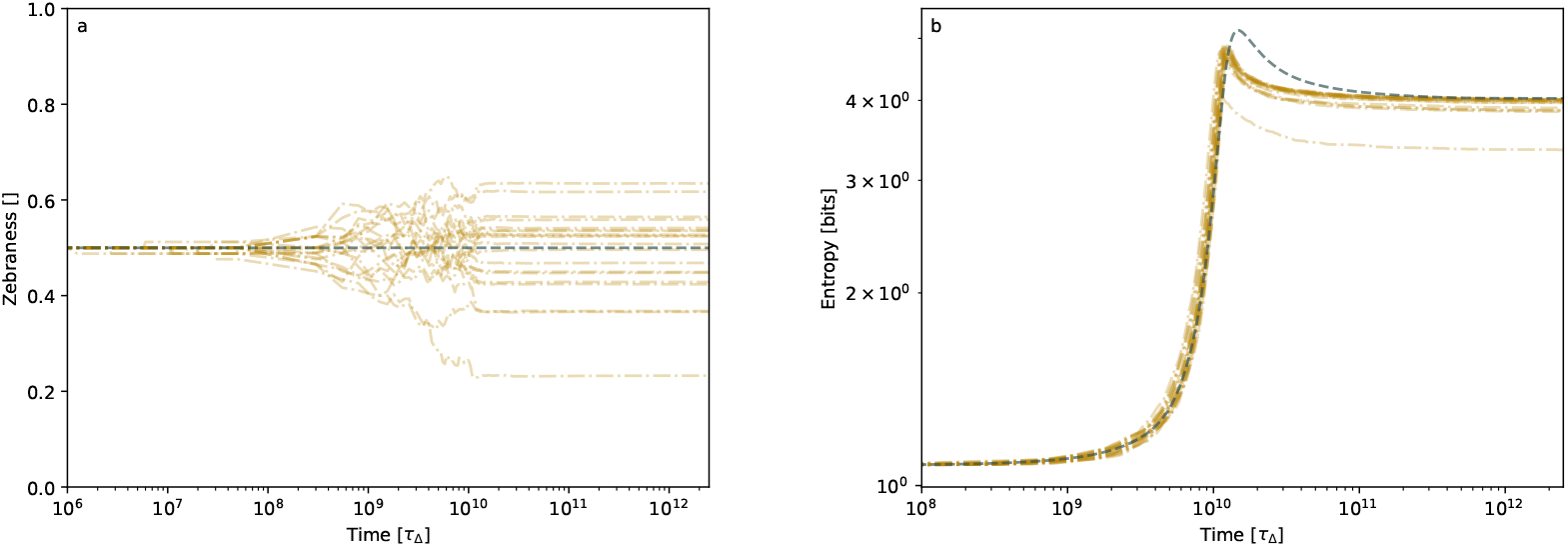
Time evolution of (a) the system-level zebraness and (b) the motif entropy, as in Fig. 4, but for parameter set 1, in which ligation stalling is activated.

A clear difference in the behavior is seen only in the system-level zebraness (Fig. 6a). As in the case without ligation stalling (Fig. 4a), the strand reactor simulation produces fluctuations during the growth phase, but the resulting bias in the zebraness then persists in the presence of ligation stalling. Ref. [9] revealed that this is due to the tendency of ligation stalling to suppress ligation in the presence of mismatches, which would otherwise counteract the bias. Hence the persisting bias in the individual strand reactor simulation trajectories corresponds to an effect of locked-in fluctuations. Fluctuations are not accounted for in the mean field description of the motif rate equations, such that their dynamics consistently stay at a zebraness of 0.5 in Fig. 6a, which is the ensemble average of the strand simulation runs.

Next, we turn to the effect of cleavage by considering parameter set 2. Cleavage drastically changes the time evolution of the mean strand length in the strand reactor simulation (Fig. 5b), creating a plateau shortly after the onset of growth, followed by a second phase of growth settling at a higher plateau at much later times. The origin of this behavior, studied in detail in Ref. [9], is visible in the time evolution of the zebraness and entropy shown in Fig. 7: The combination of ligation stalling and cleavage leads to spontaneous symmetry breaking in sequence space, where the sequence composition of the pool ultimately converges either to purely homogeneous sequences (zebraness 0) or purely alternating sequences (zebraness 1), accompanied by a drop in the sequence entropy (Fig. 7b). Both cases lead to the formation of more stable double-strands, which are less susceptible to cleavage, producing longer strands on average (Fig. 5b). The crucial dynamics are defined by the typical templated ligation time scale, setting the first plateau, and the typical cleavage time that is longer in this case and leads to the second plateau.

**FIG. 7.**
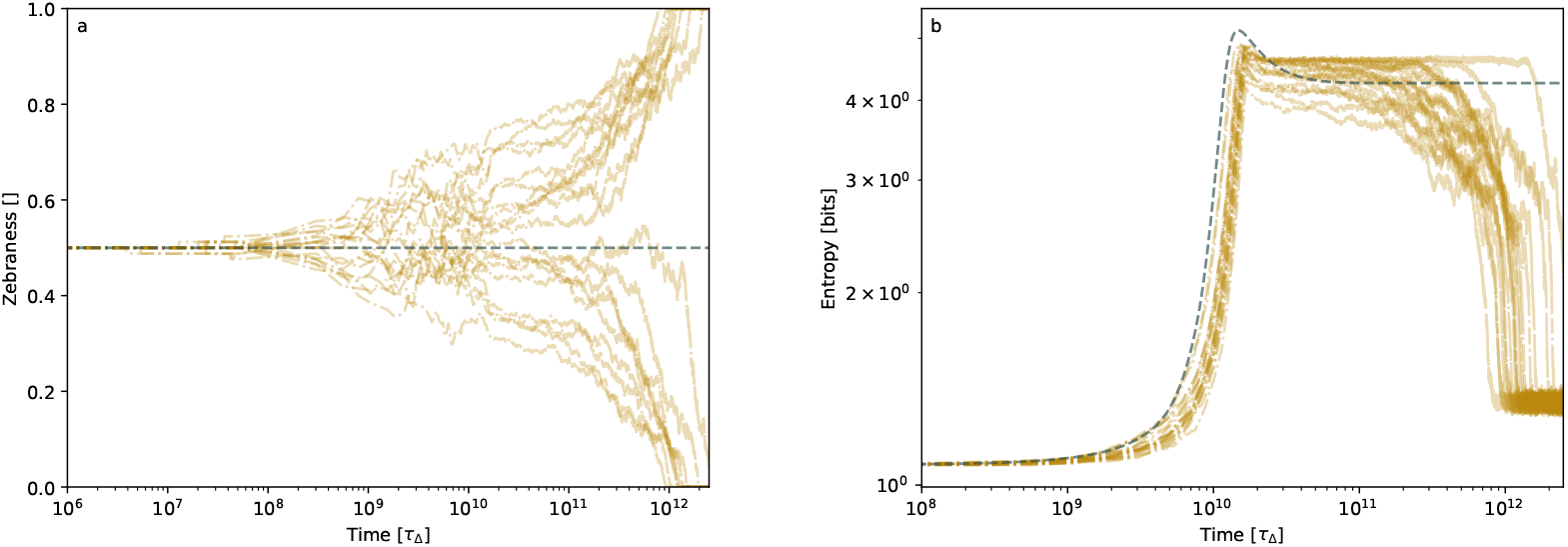
Time evolution of (a) the system-level zebraness and (b) the motif entropy, as in Fig. 4, but for parameter set 2, in which ligation stalling and cleavage are activated.

This intricate interplay between stochastic effects and hybridization-inhibited cleavage cannot be captured by the motif rate equations. It comes as no surprise then that there are significant qualitative deviations between motif rate equations and strand reactor simulation in Figs. 5b and 7. The origin of these deviations lies in the two inherent limitations of the motif rate equations already discussed above, (i) the missing effect of stochasticity within the mean field approach, and (ii) the missing kinetic effects caused by longer hybridization sites. Nevertheless, while strands are still short, up to a mean strand length of about 12 nucleotides, the motif rate equations describe the strand reactor simulation. The zebraness always remains at the unbiased value of 0.5 within the strand reactor simulation, consistent with the mean field approach that cannot account for spontaneous symmetry breaking in a finite pool. As a result, the entropy increases within the motif rate equations precisely when it decreases in the strand reactor simulation. In the strand reactor simulation, trajectories settle into one of the two zebraness regimes which reduces entrpoy, whereas the motif rate equations effectively occupy both states simultaneously, leading to a higher final entropy. The difference in the mean-length behavior at large times (Fig. 5b) can be attributed to the absence, in the motif rate equations, of the protective effect provided by stable long hybridizations against cleavage. In the strand reactor simulation, such stable hybridizations delay growth until the first plateau and protect strands from breaking after the typical cleavage time. In contrast, the motif rate equations do not account for full hybridization sites, and therefore strands continue to grow past the first plateau, eventually reaching approximately the same value as in the strand reactor simulation at very late times.

So far, we did not obtain an emerging zebraness bias within the motif rate equations, since they cannot capture the spontaneous symmetry breaking that occurs within the strand reactor simulation in parameter set 2. In principle, symmetry breaking in sequence space could emerge within the motif rate equations in two ways. First, via the energy parameters, specifically by introducing a hybridization bias as we will do next. Second, by amplification of a small bias in the initial conditions, which we investigate further below.

In both, parameter sets 3 and 4, the hybridization bias is activated, which means that the hybridization of alternating sequences is energetically more favorable, and hence more stable, than for homogeneous sequences. The hybridization bias does not have a strong effect on the time evolution of the mean strand length within the strand reactor simulation: For deactivated cleavage (parameter set 3), the hybridization bias enhances growth only slightly (Fig. 8a) compared with the case without hybridization bias (Fig. 5a). In the presence of cleavage (parameter set 4), the hybridization bias does not alter the two-step growth to a mean length of around 40 nucleotides observed already without hybridization bias (Fig. 5b vs. Fig. 8b).

**FIG. 8.**
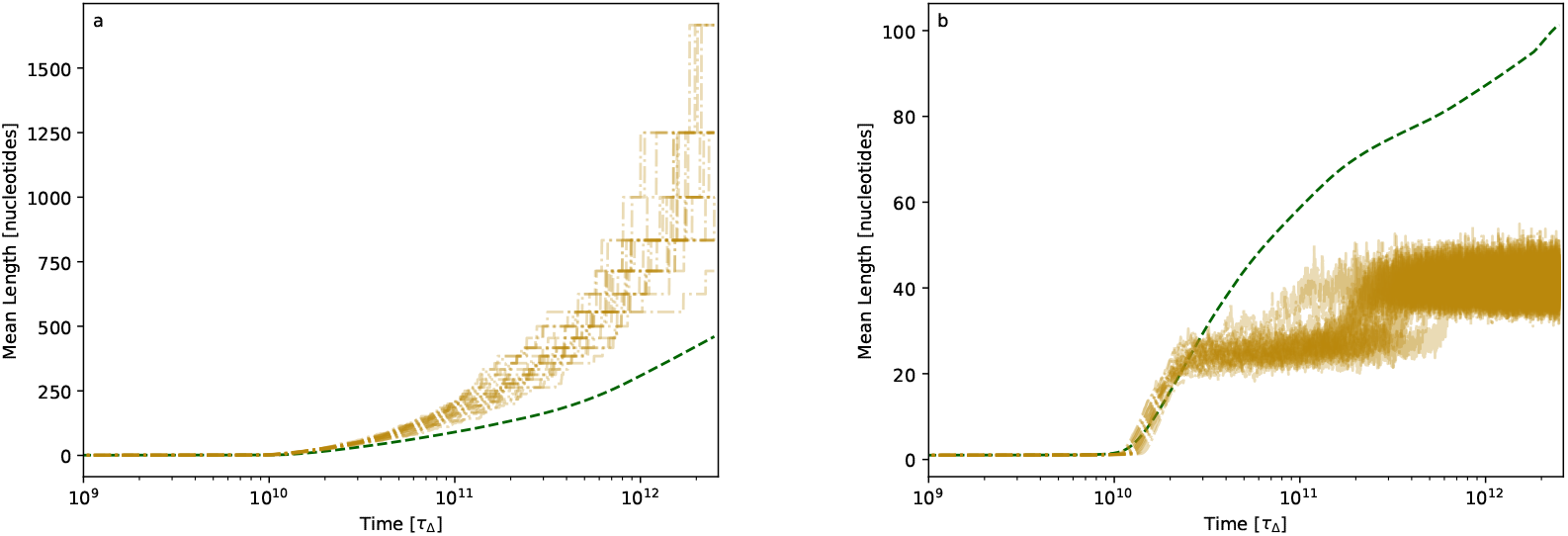
Time evolution of the mean strand length, as in Fig. 5, but for (a) parameter set 3 and parameter set 4, probing the additional effect of a hybridization bias.

In contrast, the predicted dynamics of the mean length within the motif rate equations is substantially changed by the hybridization bias. Whereas they overestimate strand growth without cleavage in the absence of the hybridization bias (Fig. 5a), they underestimate growth in the presence of the hybridization bias (Fig. 8a). With cleavage, the motif rate equations are similarly sensitive to the hybridization bias (Fig. 5b vs. Fig. 8b). This illustrates that although the estimation of the mean strand length from the motif rate equations (Appendix C 1) is quite accurate at short lengths, it is generally not robust. This comes as no surprise, since the motif rate equations focus on the informational dynamics on the scale of short motifs.

Still, the motif rate equations capture the informational dynamics for both parameter sets, 3 and 4 almost accurately, see Fig. 9, and Fig. 10, respectively. The hybridization bias leads to an emerging zebraness bias for both cases, without cleavage (Fig. 9a) and with cleavage (Fig. 10a). In the latter case, the zebraness bias even reaches its maximum value of *Z* = 1 in the long time limit, corresponding to a pool consisting only of alternating sequences. This behavior is also captured within the motif rate equations. Without cleavage, the motif rate equations reproduce the dynamics compatibly with the strand reactor simulation (Fig. 9).

**FIG. 9.**
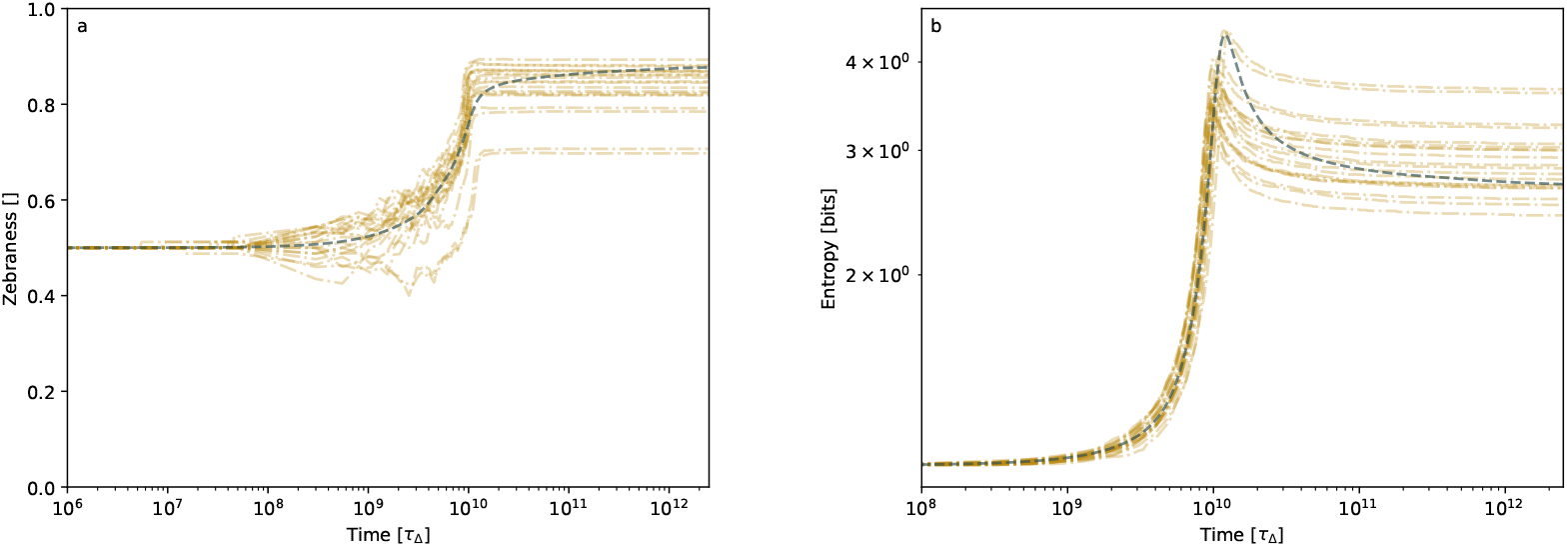
Time evolution of (a) the system-level zebraness and (b) the motif entropy, as in Fig. 6, but for parameter set 3, in which the hybridization bias is activated.

**FIG. 10.**
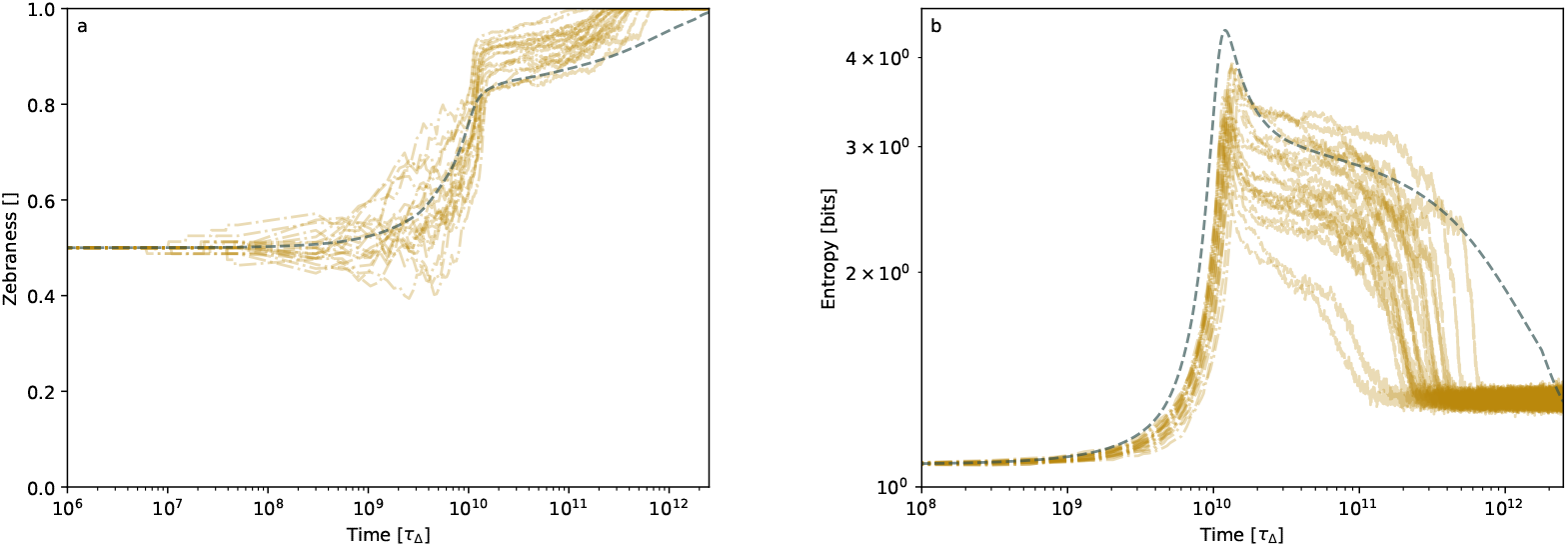
Time evolution of (a) the system-level zebraness and (b) the motif entropy, as in Fig. 7, but for parameter set 4, in which the hybridization bias is activated.

Taken together, although the motif rate equations cannot capture effects due to continuous state space size or stochasticity, they can otherwise describe the informational dynamics of the strand reactor simulation in motif space on at least semi-quantitative level. Given that the description via motif rate equations is computationally much more efficient than strand reactor simulation, this suggests that motif rate equations are a useful tool to rapidly explore the parameter dependence of the informational dynamics. This approach could be used, for instance, to identify regions within the parameter space that potentially exhibit interesting behaviors, which can then be studied in detail using strand reactor simulation. As a small illustration for this approach, we briefly consider the dependence on the initial conditions in the following.

### Probing fluctuation effects via biased initial conditions

Above, we saw that the spontaneous, noise-driven symmetry breaking in sequence space, which occurs within the strand reactor simulation for parameter set 2, cannot be described by the motif rate equations (Fig. 7). However, we can still use the deterministic description of the motif rate equations to probe whether an initial fluctuation that breaks the symmetry will grow or be attenuated, or whether the bias remains at the initial level.

Towards this end, we probe the behavior of the motif rate equations for different zebraness biases in the initial sequence pool. As before, the initial pool is dominated by monomers, with an equal number of X and Y. However, instead of the unbiased initial condition with 10 dimers of each type, we start, e.g., with 18 alternating dimers (9 each of ‘XY’ and ‘YX’) and 22 homogeneous dimers (11 each of ‘XX’ and ‘YY’). Fig. 11 shows the resulting informational dynamics obtained from the motif rate equations with parameter set 2. The plots also show the curves for any other possible combination of alternating and homogeneous dimers in the initial pool (at the same total of 40 dimers).

**FIG. 11.**
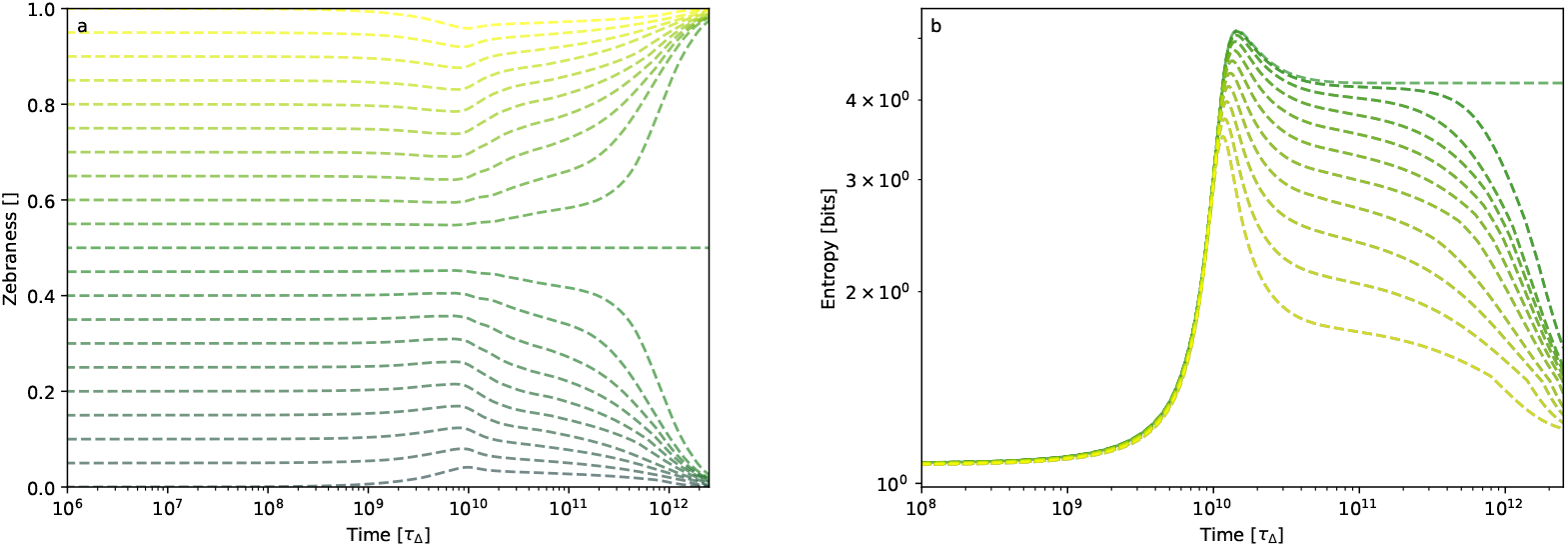
Time evolution of (a) the system-level zebraness and (b) the motif entropy within the motif rate equations for parameter set 2, starting from different initial conditions, in which the dimers within the initial pool have different biases towards alternating or homogeneous sequences.

As seen from the zebraness and motif entropy in Fig. 11, the sequence pool clearly tends to more biased states after the initial growth as expected. Mimicking fluctuation amplitudes in the stochastic system, the zebraness reliably approaches 0 or 1 in the long-time limit as observed in the strand reactor simulation. This confirms that the spontaneous symmetry breaking in the stochastic strand reactor simulation has a signature in the deterministic dynamics of the motif rate equations. Such signatures can be used for rapid parameter screens based on the motif rate equations.

## CONCLUSION AND OUTLOOK

We analyzed the informational dynamics in pools of polynucleotides, which bind to each other and react via ligation and cleavage processes. Focusing on the level of short sequence motifs, we developed a system of coupled motif rate equations to directly describe the motif dynamics extracted from full stochastic simulations on the level of polynucleotide strands, using the previously established ‘RNA reactor’ simulation framework [9, 13, 23].

Our comparison of the predictions of the motif rate equations with the strand reactor simulation was based on five different parameter sets with distinct behaviors, and on four observables: motif concentrations, motif entropy, ‘zebraness’ [9], and mean strand length. As might be expected based on the projection of the dynamics into motif space, the motif rate equations can generally not accurately predict the time evolution of the mean strand length in the strand reactor simulation. In contrast, we found that many aspects of the informational dynamics were well described by the motif rate equations. This is noteworthy, in particular, because the motif rate equations do not involve any fit parameters. Instead, all parameters of the motif rate equations are directly determined from the parameters of the RNA reactor model. We also found systematic deviations between the motif dynamics predicted by the motif rate equations and the strand reactor simulation. These could be attributed to effects depending on the influence of longer hybridization sites than the motif length, as well as stochasticity and finite system size effects.

Since the numerical solution of the motif rate equations is computationally much more affordable than the strand reactor simulation, and the motif rate equations capture many aspects of the informational dynamics on a semi-quantitative level, it will be useful to scan the parameter space for phenomena of interest, which can then be studied in more detail within the strand reactor simulation. This approach will have to be fully developed in the future. However, as a first illustration, we showed that the deterministic dynamics of the motif rate equations display a signature of the spontaneous symmetry breaking in sequence space, which occurs in the stochastic strand dynamics in a certain parameter regime. Another interesting future direction is to establish a systematic parameter inference from experimental data using the motif rate equations.

## ACKNOWLEDGEMENTS

This project was supported by the Deutsche Forschungsgemeinschaft (DFG, German Research Foundation) under Germany’s Excellence Strategy (EXC-2094-390783311, ORIGINS) and via the CRC/TRR 392 Molecular Evolution (Project ID 521256690).

## AUTHOR CONTRIBUTIONS

J.H.-K. performed the research. J.H.-K., T.A.E. and U.G. designed the research, with input from L.B. T.G. provided the code and help with the strand reactor simulations. J.H.-K., T.A.E. and U.G. wrote the paper, with input from all authors.

## Appendix A

**Motif Rate Equation Terms**

In the following, we discuss the individual terms of the motif rate equations in detail. A test of the implementation is done in Appendix B where we compare the numerical solution to the exact solution of a simple monomer-dimer system that can be solved analytically. All implementations in this paper use the Motif Reactor Simulation, Analysis and Inference Kit in Python (MoRSAIK) [10].

Let ℳ_ℓ_ be the set of all sequence motifs with a length of ℓ nucleotides. Mathematically, the motif rate equation terms depend on the motif concentrations that we store in the motif concentration vector 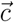, element of the *motif concentration space* 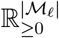 with semi-positive real values and dimension of the number of motifs equal to the cardinality of the motif set | ℳ_ℓ_|. Every motif rate equation term indicated by an index *i* maps between identical motif concentration spaces: 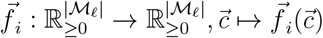.

### 1. Motif Production Rate Calculation

The motif production rate becomes zero if the reaction is not allowed by our convention, i. e. if the second nucleotide of one reactant or the product is empty (zero), or in case of an empty spot between two nucleotides. If our convention is fulfilled, the motif production rate of a specific reaction is given by the product of the corresponding motif production rate constant multiplied by the concentrations of the reactants and template, as in Equation (6). This implies that we need to introduce an exceptional equation in case the left strand is a monomer, *l* = (0, *l*_2_, 0, 0) (and *r* = (0, *r*_2_, *r*_3_, *r*_4_)). In this case, there is no left product, as in Equation (3), and the central product becomes

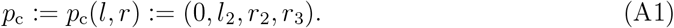

In case the right reacting strand is a monomer, Equation (4) still holds, but there is no right produced motif as in Equation (5).

The boundaries for the sums in the motif production terms for produced motifs follow from our convention. Accordingly, the first and last nucleotides of each motif can be empty or a letter. Consider the left reactant motif, *l* = (*l*_1_, *l*_2_, *l*_3_, *l*_4_), right reactant motif (*r* = (*r*_1_, *r*_2_, *r*_3_, *r*_4_)) and the template motif *t* = (*t*_1_, *t*_2_, *t*_3_, *t*_4_), then *l*_1_, *r*_4_, *t*_1_, *t*_4_ ∈ 𝒜_0_. The second nucleotide, as well as the third nucleotide of the template, is always a letter, *l*_2_, *r*_2_, *t*_2_, *t*_3_ ∈ 𝒜. The third nucleotide of each motif is only empty if the motif is a monomer, i. e. the first and the fourth nucleotide are empty, *l*_3_ ∈ 𝒜 ∪ (*{l*_1_*}* ∩ *{l*_4_*}*), *r*_3_ ∈ 𝒜 ∪ (*{r*_1_*}* ∩ *{r*_4_*}*).

The motif production rate constants are approximated by the extension rate constants discussed next. To ensure positivity, we use logconcentrations to compute the motif production terms, the details of their computation are discussed in Appendix A 3.

### 2. Extension Rate Constants

We approximate the motif production rate constants by extension rate constants *k*_ext_ assuming that the motifs fully capture the participating strands. Thus, a produced four-nucleotide motif is assumed to be a tetramer, a beginning or ending motif a trimer, etc.. As in [9], the extension rate constants are given by the ligation rate constant (*k*_lig_) divided by the dissociation constant of the complex *C* resulting from the hybridization of produced and templating motif,

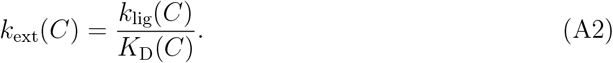

The ligation rate constant is given by a sequence independent ligation rate constant 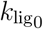 for perfect matches multiplied by the stalling factors Φ_*±*_.

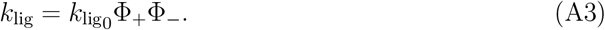

The stalling factors are the product of the stalling parameters for mismatches close to the ligation spot.

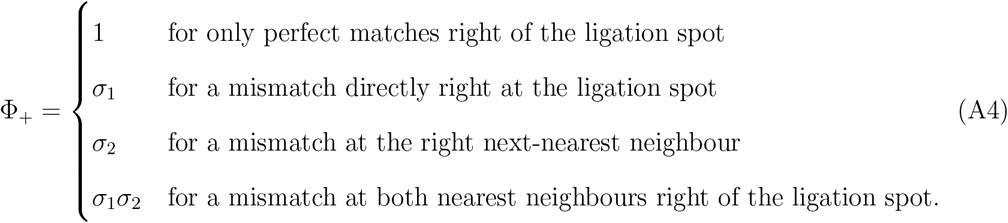

Φ_−_ analogously for the mismatches left of the ligation spot.

The dissociation constant is calculated from the hybridization energy difference, as defined in Equation (1). The hybridization energy is given by the sum of hybridization energies of the two by two nucleotide segments in the complex, where dangling ends have a hybridization energy of *ϵ*_com_ *± δ*_*ϵ*_*/*2 if they are perfectly matching and their sequence is alternating (+) or homogeneous (−), and *ϵ*_1nc_ if there is one mismatch, fully hybridized segments have a hybridization energy of 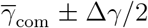if they are perfectly matching and their sequence is alternating (+) or homogeneous (−), and *γ*_1nc_ (*γ*_2nc_) if they have one (two) mismatches. The values of the reaction parameters are taken from [9] and stated in Table V.

### 3. Motif Rate Equations with Log-Concentrations

To ensure that the concentrations stay non-negative, we express the differential equations in terms of the *log-concentrations* 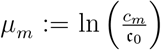 and choose very low concentrations for non-occupied motifs. The corresponding vector is 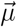 with log-concentrations of each corresponding motif as entries 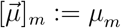. The motif rate equation is given by

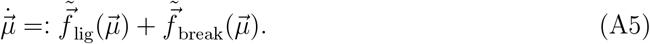

The individual terms transform to

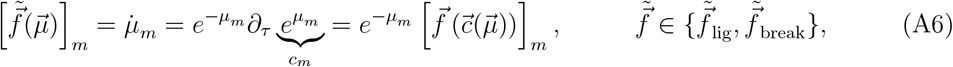

where 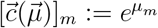.

The total motif production rate in terms of log-concentrations then becomes

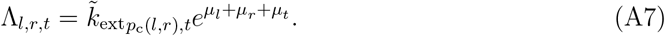

As before, we split the motif production terms into terms for the left reactant 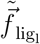, the right reactant 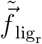, and the three resulting motifs 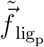. Both, the ending as well as the leaving strand, are consumed in this reaction. This can be expressed in terms of log concentrations.

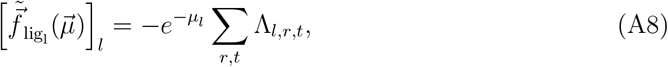

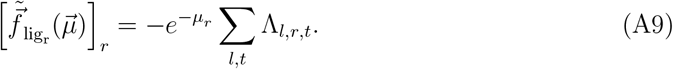

For the three produced motifs (*p*_l_, *p*_c_, *p*_r_), we get

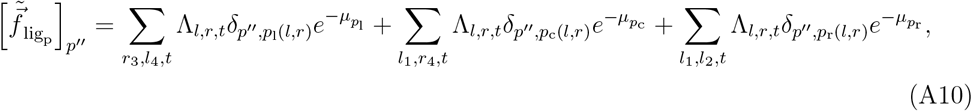

with the exceptional case of monomers, which do not produce left (or right) products according to our convention, see Appendix A 1.

The breakage terms for log-concentrations split into terms for breaking 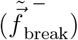 and resulting 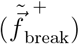 motifs,

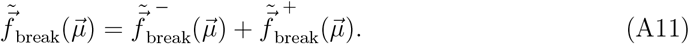

The terms for breaking motifs are

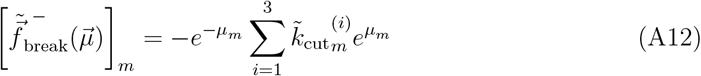

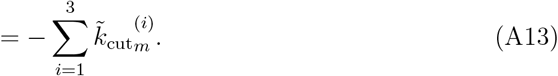

The terms of the resulting motifs

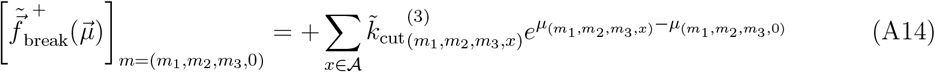

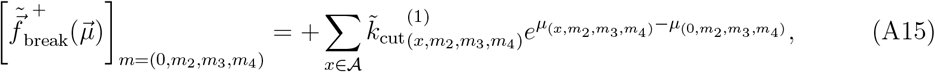

with all other entries being zero.

Summing all contributions together gives the overall motif rate equations for logconcentrations.

### 4. Smooth Cutoff

To model finite size effects in the mean field approach of the motif rate equations, we introduce a cutoff *δ*_0_ for the reaction rates: if the concentration of any reactant falls below this cutoff, the corresponding reaction rate is set to zero [1, 3, 12, 28]. To prevent sharp discontinuities caused by this clipping, we further introduce a threshold *δ*_1_ and a smooth cutoff function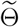. This function evaluates to zero when the concentration is below *δ*_0_ and to one when it exceeds the threshold, with a smooth cosine-based transition between these two values for intermediate concentrations.

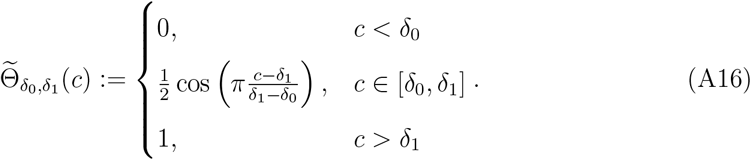

We then apply this smooth cutoff to the reaction rates:

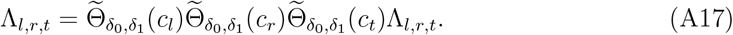

As a physical interpretation, the threshold acts as a pseudo-count, and analogously, the cutoff can be viewed as a pseudo-zero. In this work, we set the threshold to the concentration of a single particle (c_Δ_) and the pseudo-zero to half of that value.

### 5. Mass Conservation

In a scenario without influx or outflux, nucleotides are not lost; they are merely rearranged among motifs. Consequently, mass is conserved in the motif rate equations discussed in the Methods Section and in Appendix A. To maintain mass conservation and compensate for numerical inaccuracies, we introduce mass-conservation rates that renormalize via forcing. These rates effectively simulate a harmonic potential with its minimum at the total initial mass, while ensuring that the number of beginnings equals the number of ends such that every strand retains exactly one beginning and one end.

#### Total Mass

Computing the total mass is nontrivial, as a single nucleotide can be part of several motifs, and only the concentration of the latter are followed by the simulation. Let us call the *total mass M* of the system the concentration of nucleotides in the system and the mass *M*_*a*_ of a specific nucleotide *a* ∈ 𝒜 the total concentration of nucleotides *a*. The total mass can be computed by the sum over the mass of all nucleotides. We compute the nucleotide specific mass by scanning all motifs for the specific letter and adding the mass accordingly. There, we have to carefully consider our convention to prevent double counting.

According to our convention, the second nucleotide spot of every motif is occupied. It always fully contributes to the specific mass. The first and the last nucleotides never contribute to the mass, since they are tracked in the motifs before or after the current motif. The third letter only contributes, if the last nucleotide is empty, since the motif that would follow does not fulfill our convention and therefore is not tracked.

Thus, we have the following expression for the different nucleotide masses:

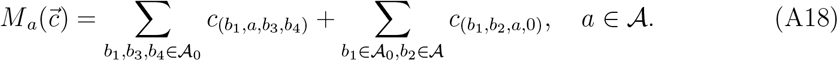

Those are separately conserved for zero influx and outflux.

Additionally, we demand that every strand needs to have a leaving and an ending strand. Otherwise, it would be infinitely long. This requires that the concentration of the beginning motifs equals that of ending motifs, and therefore

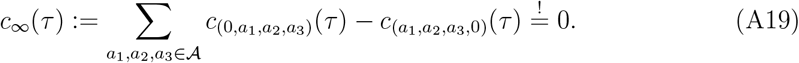

#### Mass Conservation Rates

To enhance numerical stability when solving the motif rate equations, we enforce consistent mass conservation using a harmonic potential as a loss function. This potential is minimized when total mass is conserved and when the number of beginning motifs equals the number of end motifs. It consists of two terms: First, a mass-conservation term, minimized when total mass is preserved and weighted by the mass conservation rate constant *k*_mass_; and second, a strand-completion term, minimized when the concentration vector contains equal numbers of beginning and end motifs, weighted by the strand-completion rate constant *k*_strand_.

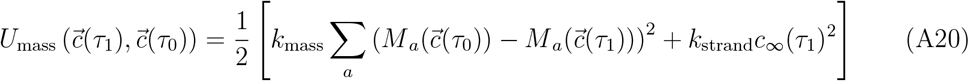

The gradient of this mass-conservation potential yields the correction terms that we add to the motif rate equations:

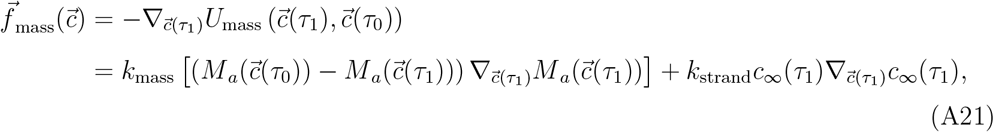

with

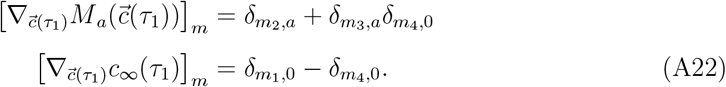

For the purposes of this paper, we have set both, the mass conservation rate constant and the strand completion rate constant to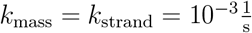.

## Appendix B

**Monomer-Dimer System**

In the following, we test the motif rate equation by comparing the integrated solution to the analytic solution of a simple monomer-dimer system with a single-letter alphabet.

Let *m*_1_ denote a monomer sequence, *m*_2_ a dimer. The total mass of the system — i. e., the total number of nucleotides *M* — is therefore

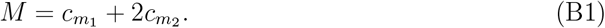

From this, the dimer concentration can be expressed in terms of the monomer concentrations, reducing the motif rate equations to a single equation:

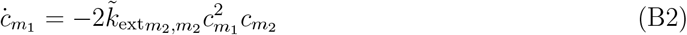

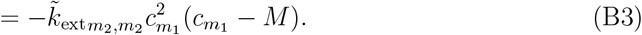

We introduce the *monomer frequency* 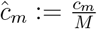, whose time evolution is given by

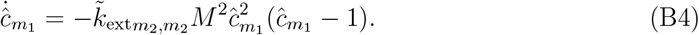

Integrating yields

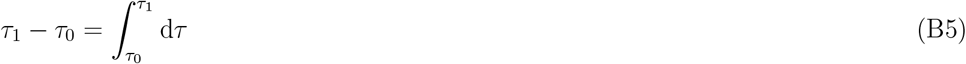

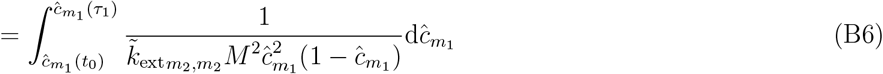

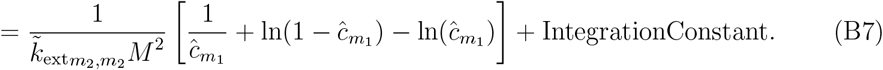

The IntegrationConstant is determined by the initial conditions at 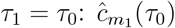,

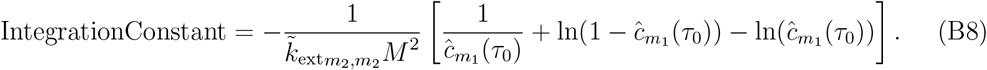

This provides the full solution:

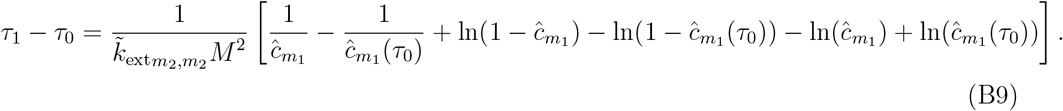

This case is equivalent to solving the motif rate equations with motif production rate constants 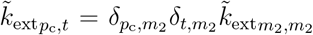. The resulting motif concentration trajectories exactly match the numerical solution of the motif rate equations(see Figure 12) validating our implementation.

**FIG. 12.**
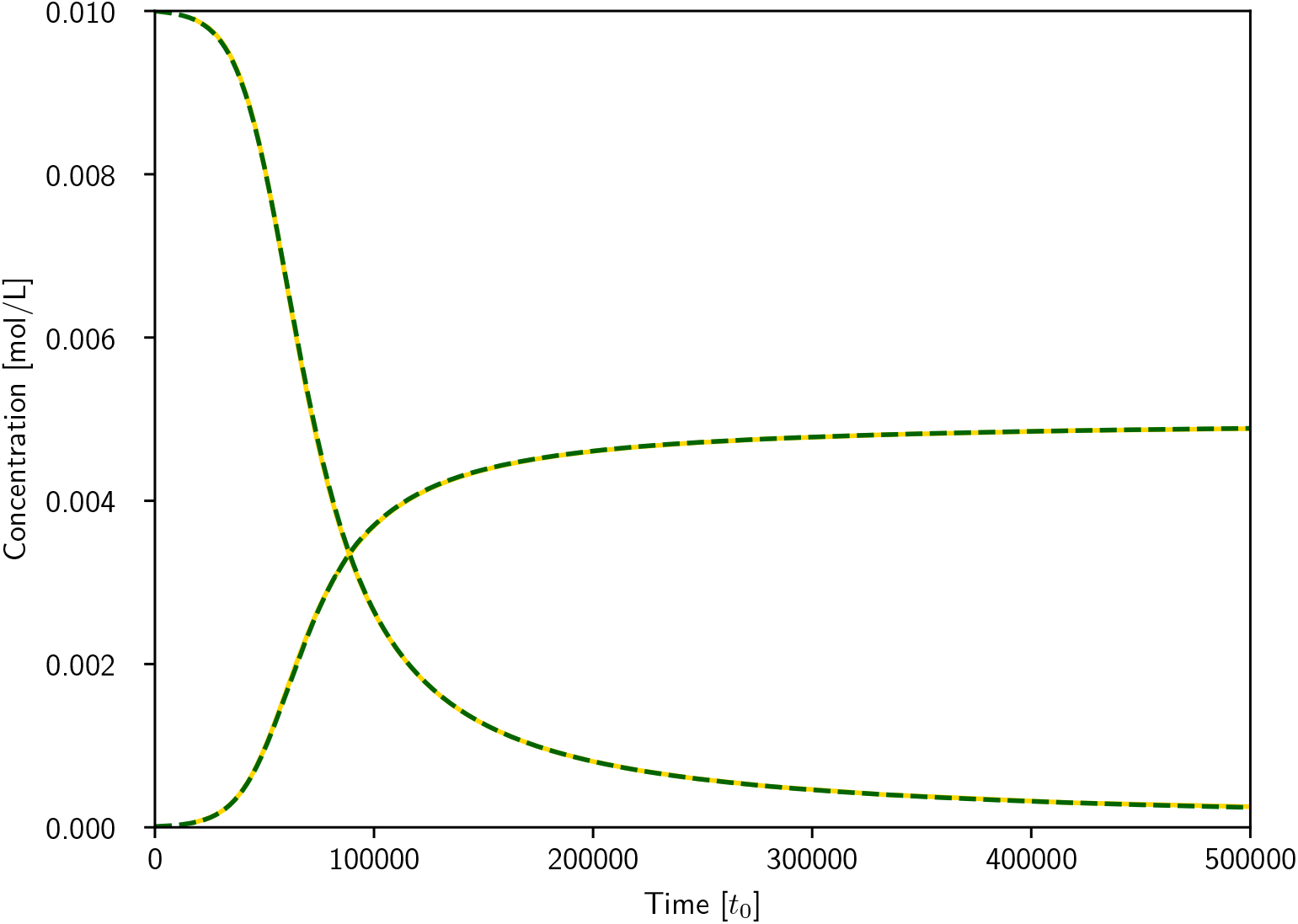
Monomer (decreasing) and dimer (increasing) concentration trajectories determined analytically (solid gold) and numerically (dashed green).

## Appendix C

**Observables**

### 1. Mean Strand Length

While we can explicitly determine the mean strand length in the strand reactor simulation by averaging over the strand lengths from the output of the simulation, we need an estimate of it for the motif rate equations. For this, the length of all explicitly captured strands – the monomers and dimers – contribute directly. The mean length of longer strands needs to be estimated from the concentration of beginnings, continuations and ends. Beginnings contribute by one nucleotide to the mean length as their third and fourth nucleotide is considered by corresponding continuations or ends. For the same reasons, also continuations contribute to the mean length with one nucleotide. Ends contribute to the mean length with two nucleotides, since the second and the third nucleotide are not considered by other motifs. The total number of strands, 𝒩_strands_, is equal to the number of explicitly captured strands and ends [34].

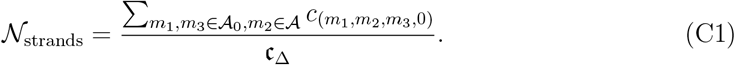

By taking the ends and considering that explicitly captured strands also have an empty spot at the end, the *mean strand length* 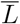 reduces to

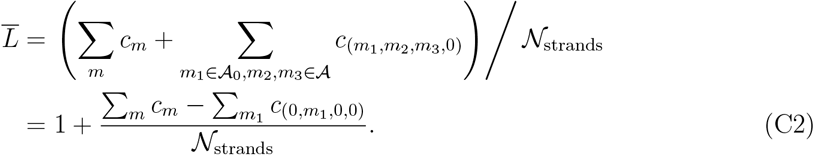

### 2. Onset of growth in a Finite System

Here, we will estimate the difference of the onset of growth for the motif rate equations and the strand reactor simulation due to different system sizes.

Göppel et al. [9] derived an estimate for the onset of growth

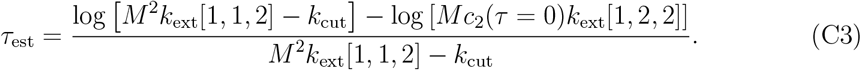

given the total number of nucleotides *M* and the sequence averaged extension rate constants,

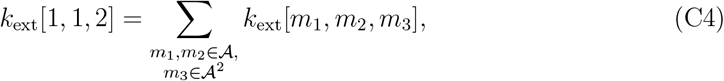

and approximating that the dimer concentration *c*_2_ grows exponentially in the initial growth phase,

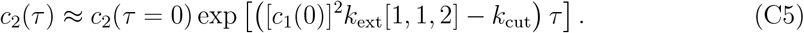

In a finite system, the onset of growth might be delayed, if the first dimer is produced after *τ*_est_. Let c_Δ_ be the concentration of a single strand and *τ*_dim_, the time at which the first dimer is produced. We use the exponential growth approximation for the initial dimer concentrations,

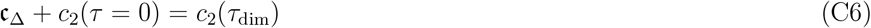

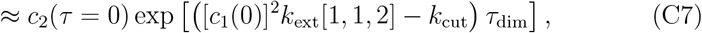

and solve for *τ*_dim_,

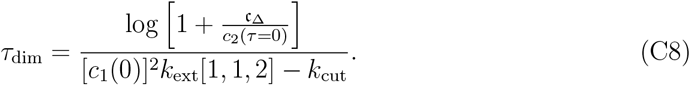

If the initial pool consists mainly of monomers, the onset of growth gets delayed, if

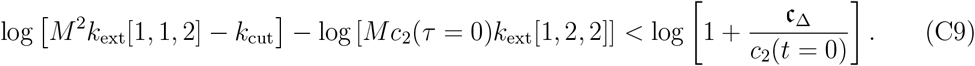

Solving for c_Δ_ gives

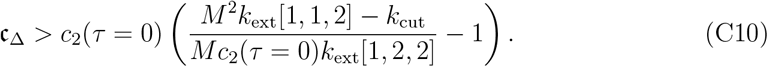

For parameter set 0, the onset of growth is at 3.3 *·* 10^9^*τ*_Δ_, while the first dimer is produced before 2 *·* 10^7^*τ*_Δ_. Thus, the onset of growth is not delayed due to the finite size.

